# Towards Video-LLM Driven Workflow for Behavioral Segmentation and Scoring in Mice Performing a Skilled Water Reaching Task: An Evaluation of Recent LLM Models

**DOI:** 10.64898/2025.12.12.694037

**Authors:** Tony Fong, Hao Hu, Haozong Zeng, Timothy H. Murphy

## Abstract

**Significance:** Behavior scoring is labor-intensive and subjective, introducing variability in results. Large Language Models (LLMs) capable of video understanding offer a transformative solution to manual scoring, crucial for accelerating and standardizing neuroscience workflows.

**Aim:** We sought to benchmark state-of-the-art video LLMs (Gemini 2.5 Pro, Qwen3-VL, and VideoLLaMA3) for automated behavioural segmentation and scoring of mice performing a water-reaching task.

**Approach:** Videos of mice performing water reaching from the front view were analysed by the LLMs. Accuracy was compared across different models and against prompt adjustments within Gemini. To assess classification determinants, video fidelity was altered through pixel interpolation and key regions blurred (paws/snout-mouth). In addition, the models were asked to describe the mouse’s actions over time.

**Results:** Gemini 2.5 Pro (0.74 ± 0.12 accuracy) and Qwen3-VL-30B (0.67 ± 0.13) exhibited ability to classify trial outcomes. Reliable classification required a minimum pixel resolution of 0.28 mm per pixel. Accuracy is significantly reduced upon obscuring the snout-mouth area. In 549/1058 of videos, Gemini 2.5 Pro also provided completely accurate frame-to-frame behaviour segmentations.

**Conclusions:** Video-LLMs offer potential to accelerate neuroscience by providing scalable, objective quantification of goal-directed behaviors. By producing temporal annotations, Gemini enables fast first-pass labelling that markedly streamlines manual dataset curation.

## 1. Introduction

In neuroscience, behavior remains the primary outcome of interest across many diverse subfields. As such, quantitative and unbiased assessments of animal behavior is critical, yet traditional scoring methods are not only laborious, but also prone to human errors and biased by subjective interpretation even with corrective efforts (1–3). Over the past decade, computer vision methods—most notably convolutional neural network (CNN)-based frameworks such as DeepLabCut, SLEAP, and lighting pose—have revolutionized behavioral studies by enabling markerless pose estimation in 2D and 3D (4–9). When combined with clustering algorithms that group keypoints into behaviorally meaningful states, these methods have enabled supervised and unsupervised classification of specific behaviors (10–15). These approaches represent an important step toward standardization and reproducibility in behavioral neuroscience. However, pose-based clustering alone remains insufficient for understanding how animals interact with the environment—particularly in trial-based tasks where outcomes (e.g., hits versus misses) are critical. Although hardware sensors, such as capacitive sensors (16,17), IR beam breaks (18), and motor encoders (19) can aid in determining trial outcomes, they constrain experimental design and limit the range of tasks that can be tested. In particular, skilled forelimb reach-to-grasp and retrieval tasks provide sensitive measures of rodent motor circuitry, yet their scalability can be hindered by time-consuming and subjective manual outcome scoring (20–23). At present, there are no robust out of the box video-based approaches that do not require fine tuning that can distinguish successful from failed reaching attempts.

In recent years, the rapid evolution of large multimodal models (LMMs) has opened new frontiers in video understanding (24). Many LLM-based models, such as VideoPrism (25) from Google DeepMind and MouseGPT (26) can achieve human-level performance in pose estimation and self-directed behaviors classification from raw videos alone, reaching parity with expert scorers on benchmark datasets of mouse videos. These advances suggest that general-purpose video LLMs trained on large and diverse datasets may extend beyond recognizing self-directed behaviors to also classifying how animals interact with—and produce outcomes in—their environment. Most recently, the release of Gemini 2.5Pro (27), and Qwen3-VL (28,29) models surpass prior models across a range of video understanding benchmarks, raises the possibility that video LLMs could generalize to mouse behavior scoring without additional training.

Here, we introduce a workflow that leverages video LLMs to directly segment and score rodent reach-to-grasp behaviors from single-view videos. This approach lays the foundation for providing an efficient and scalable alternative to traditional pipelines that rely on pose estimation, feature engineering, and clustering. Using this framework, we benchmark several recent video LLMs on headfixed mice performing a water-reaching task. Our results suggest that video LLMs may streamline behavior analysis, reduce hardware and annotation overhead, and offer fully generalizable scoring capabilities for behavioral neuroscience.

## Methods

### Animals and Surgery

Male C57BL/6 mice, five to six months of age and of varied genotypes, were used in this study. Animals were maintained under a standard 12/12 h light/dark cycle (lights on at 7:00 A.M.). Mice were implanted with a chronic transcranial window and a head-fixation bar as previously described (30). All experimental procedures were approved by the University of British Columbia Animal Care Committee and were conducted in accordance with national guidelines.

### Water Reaching Training and Testing

Following post-surgical recovery, mice were water restricted and trained on a custom head-fixation platform (Fig. 1A). Each session consisted of 120 trials, including 102 potentially rewarded trials (85%) and 18 null trials (15%). Approximately 10µl of water reward was delivered through a spout positioned 7 mm to the right of the animal’s snout (vertical offset ± 2 mm) controlled by a raspberry pi 4B GPIO using custom scripts. Mice underwent a minimum of 3 weeks of training prior to testing. After 7 days of baseline testing, a photothrombotic stroke was induced, followed by a 3-day recovery period. Testing then resumed once daily for an additional 24 days. Behavior capture was conducted using a global shutter raspberry pi camera (IMX296 monochrome Innomaker) recorded at a resolution of 1200 x 800 at 54 fps with 5 ms exposure under IR light to capture without motion blur.

**Figure 1.**
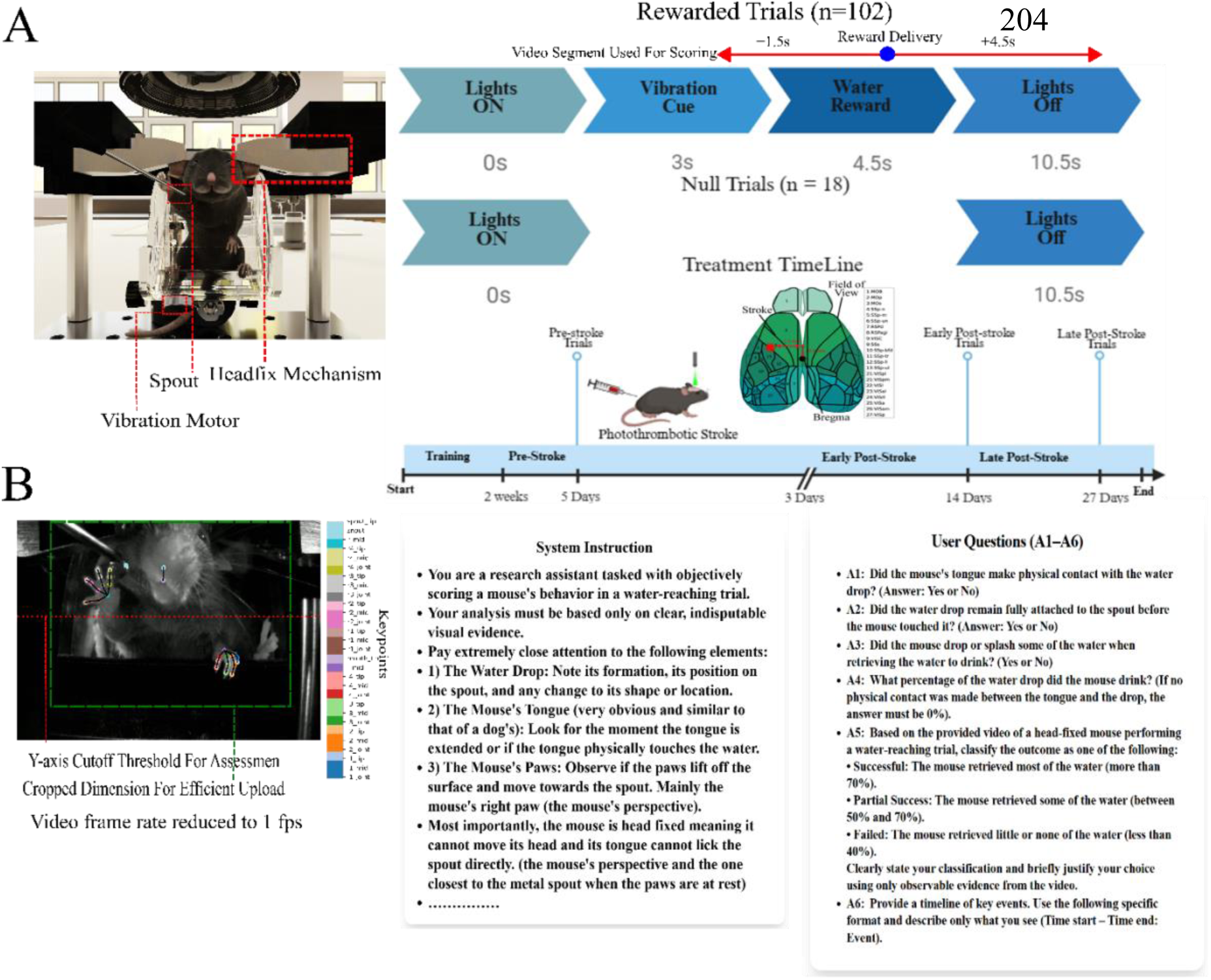
Experimental Setup and LLM input. **A.** Mouse water-reaching task and experimental timeline. Mice were headfixed in a custom setup in a standing position. A vibration motor was situated on the cup. An animal underwent 102 rewarded trials and 18 non-rewarded trials in each session. A vibration reward was delivered 3 seconds after a trial start followed by a water reward 1.5 seconds later. Six seconds after the reward the trial ends. Results of the trial are scored on -1.5s and 4.5s after the reward is delivered. Mice trained for a minimum of 2 weeks prior to testing and received stroke induction (1.5; 0.5) mm from bregma. **B.** Video data configuration with system instructions and applied prompts. The original video was cropped for efficient transfer to Gemini. Only trials where the mouse’s right paw, tracked by DeepLabCut, rises above a certain threshold are scored. All video LLMs received system instruction and user questions with the video input.

### Photothrombotic Stroke

Focal ischemia was induced by photothrombotic occlusion at a target region between the sensory and motor cortex (stereotaxic coordinates: 1.5 mm lateral, 0.5 mm anterior to bregma; Fig. 1A) as previously described (31). Mice received an intraperitoneal injection of the photosensitive dye Rose Bengal (RB; 0.1 ml/10 g body weight; R3877-5G, Sigma-Aldrich). Two minutes later, a 40mW diode-pumped solid-state 532 nm laser, attenuated to 17mW with a polarizer, was applied to the target region to induce focal ischemia. The laser beam diameter was 0.9 mm at full width at half maximum (FWHM). Prior work has shown that this procedure produces tissue damage restricted to the irradiated area.

### Videos Processing Pre-LLM Analysis

As shown in Fig. 1B–C, only the 1.5 s period before and 4.5 s after reward delivery from each video was selected for scoring. All digits from both paws, snout and mouth tip were tracked using DeepLabCut (7). Only videos in which the center of the reaching paw (average of all digits for the right paw) moved above a manually defined y-axis threshold were included in the analysis. To optimize upload speed, the videos were cropped to a resolution of 720 × 740 pixels.

### Video Manipulations

To test the effect of temporal resolution needed, videos were also re-encoded frame by frame using opencv-python (v4.8.0.76) to specific frame rates of 1, 2, 6, 13, 27, and 54 fps. Gemini would only sample/tokenize one frame per second of video playtime by default which translates to the number of frames it samples as 1/fps, i.e. the slower the frame rate the more frames Gemini would sample. However, to ensure all frames would be tokenized by the tested video LLM (Gemini, Qwen, and VideoLLama), all analysis was conducted on videos set to 1 fps unless otherwise indicated. For testing changes in video resolution, each clip was first down sampled and then up sampled back to the original dimensions prior to model input to maintain a consistent total token count. This procedure reduces the effective video resolution because fine-grained spatial details lost during down sampling cannot be recovered through up sampling, resulting in a visually smooth but information-reduced video that preserves size but not detail. To evaluate the importance of localized visual cues, the centers of the right and left paws were blurred using a Gaussian blur (σ = 0, radius= 140 pixels, kernel size = 51). The center of the mouse face (estimated as the average pixel position between the snout and mouth tip) was blurred using a Gaussian blur (σ = 0, radius = 180 pixels, kernel size = 51).

### Video LLM Analysis

Prompts used in the analysis were generated by the LLM itself using descriptive guidance by one of the authors, and the overall process for prompt generation, i.e. system instructions and user questions as shown in Fig. 2A. Only the system instructions were modified in each iteration while user questions were adjusted solely for consistency of flow. System instructions were refined in Google AI Studio (https://aistudio.google.com). The full system instructions applied can be found in Fig S1. The model and human expert results classified trials as success (visually consumed >70% of the water reward), partial success (50–70% water consumption), and failure (<40% of water consumed), as defined in the prompt (Fig. 2A). For analysis, partial and full successes (for water drop consumption per trial) were combined into a single success category to simplify scoring and comparisons. For Gemini 2.5Pro, video analysis (i.e., uploading the system instruction prompt and videos) was performed using the Google Gen AI SDK (https://github.com/googleapis/python-genai). Detailed documentation can be found at https://ai.google.dev/gemini-api/docs. A custom scheme was also created and uploaded for structured outputs from Gemini. All other open-source video LLM models were run in Python using the transformers (v4.57.1) package on a workstation equipped with a GPU featuring 96 GB of video memory (RTX-PRO-6000). Weights for Qwen3-VL-30B, Qwen3-VL-8B, and VideoLamma3 were retrieved from Hugging Face with the following links Qwen/Qwen3-VL-30B-A3B-Instruct, Qwen/Qwen3-VL-30B-A3B-Instruct-FP8, Qwen/Qwen3-VL-8B-Instruct, and DAMO-NLP-SG/VideoLLaMA3-7B, respectively. In the Qwen and VideoLLamMa models, system instructions were inputted as past chat history. Detailed instructions for employing these open source models can be found at their respective github page: Qwen (https://github.com/QwenLM/Qwen3-VL) and VideoLLaMA (https://github.com/DAMO-NLP-SG/VideoLLaMA3) both Alibaba associated repository https://github.com/QwenLM and https://github.com/DAMO-NLP-SG, respectively. All videos were analyzed with the temperature set to 0 (or 0.1 if 0 is not a valid option) to maintain consistency and top-k to 0.95 to minimize hallucinated outputs. Specific system instruction can be found in supplementary materials (Fig S1).

**Fig 2.**
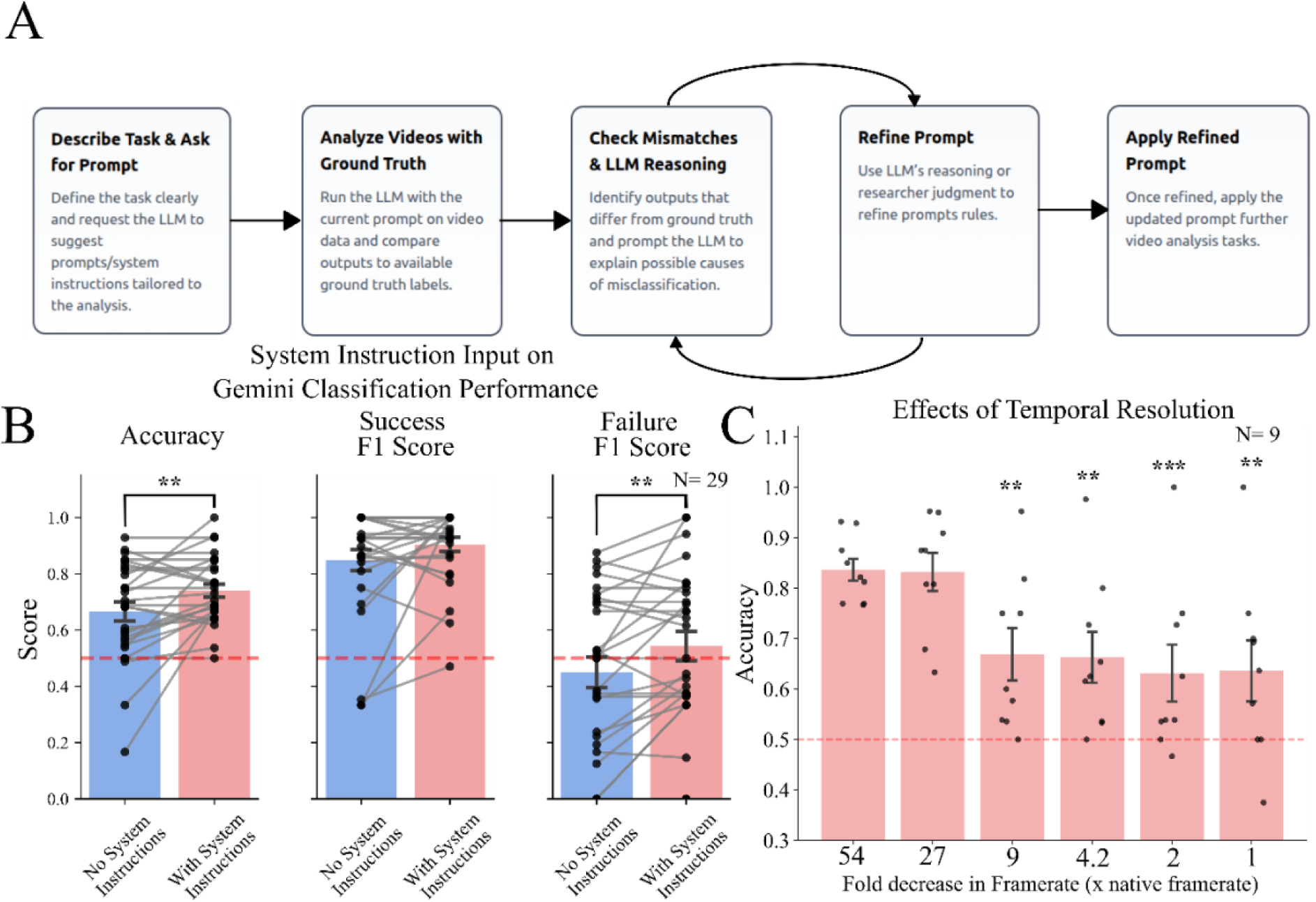
LLM Workflow and Parameters Testing. **A.** LLM prompt generation and refinement for behavioral video analysis. Only the system instructions of the prompt were continuously refined during each interaction. For simplicity, system instructions were refined in Google AI Studio. **B.** Improved accuracy of LLM with system instructions/ prompt refinement and increased temporal. The same sessions (with 6-88 videos each) are analyzed with or without system instructions added. **C.** Videos were set to a given apparent frame rate (fold decrease compared to native framerate of 54fps) in opencv prior to delivery to Gemini. The lower the frame rate the more frames the LLM would sample. The red dashes line represents the performance of a random classifier. *p < 0.05, **p < 0.01, and ***p < 0.001. Each individual point represents a session composed of 6-88 trials.

### Gemini 2.5 Pro Water Drop Quantification

To assess Gemini 2.5 Pro’s quantification ability, real still images of water drops of varied volumes (20μl, 40μl, 80μl, and 160μl) on a parafilm sheet were presented to the model. The model was instructed to detect both the number of water drops and their relative sizes, giving more weight to the smallest discernible drops, to evaluate its capacity for recognizing relative amounts.

To further test the model’s evaluation of temporal changes in quantity, videos of a single 80μl water drop undergoing volumetric manipulation were presented (using a pipette). In these videos (recorded at 30 fps), the drop’s volume was changed by +25%, 0, - 25%, - 50%, -75%, and -100% of its initial volume of 80. The model was then queried on the estimated percentage of water removed from the original 80 μl volume.

### Statistical analysis

Data are all presented as mean ±std. Statistical significance was determined using a post hoc two-way ANOVA followed by paired Student’s t tests (with Tukey’s correction) as appropriate in python. Single descriptor events are modelled as Bernoulli/ Binomial processes to calculate mean and 99% confidence intervals. Cohen’s kappa (κ) was calculated as κ = (observed agreement - expected agreement by chance) / (1 - expected agreement by chance), measuring inter-rater reliability beyond chance agreement. Values were interpreted using Landis and Koch (1977) benchmarks: 0.41–0.60 fair agreement, 0.61-0.80 as substantial agreement and 0.81-1.00 as almost perfect agreement. Prior to calculating accuracy and F1-scores, data were balanced by randomly down sampling to the minority class size within each session to ensure equal representations when benchmarking. For example, if a session with only 20 failed trials and 50 successful trials, the dataset is balanced by selecting a random subsample of 20 successful trials (and all failed trials) for the analysis. All performance metrics were calculated within each session, containing 6-88 trials. The level of significance is denoted on the figures as follows: *p < 0.05, **p < 0.01, and ***p < 0.001.

## Code and Data Availability

All codes for video LLM and statistical analysis can be found at the following github repository: https://github.com/tf4ong/videollm. All data are available at the open science frame-work repository: https://osf.io/24euy. LLMs such as Claude and Gemini were used in the assistance of code generation for statistical testing and visualization.

## Results

Mice were trained for a minimum of two weeks on a head-fixed water-reaching task. During training, the water spout was positioned approximately 6 mm lateral to the right, 2 mm ventral, and 5 mm anterior relative to the mouse’s snout. Each rewarded trial began with a vibration cue, followed by water delivery 1.5 s later. Videos were scored from 1.5 s before to 4.5 s after reward delivery. Non-rewarded trials were not scored as illustrated in Fig 1A. To optimize performance, we implemented an iterative workflow for prompt engineering (Fig. 2A). Prompts provided to Gemini consisted of two components: system instructions and user queries. The system instructions served as predefined guidelines that directed the video LLM’s reasoning process and shaped its output. We iteratively tested Gemini’s responses on ten trial videos with known ground truth (human) labels, identified incorrect outputs, and refined the system instructions accordingly. Each iteration incorporated corrections and model feedback until the generated outputs aligned with the ground truth. A paired t-test revealed that the presence of system instructions affected the accuracy (0.07 increase; p<0.01), F1 scores for failure classification (0.09 increase; p<0.01), but not success classification (0.06 increase; p = 0.1) of Gemini (Fig. 2B).

Next, we investigated the effect of temporal resolution on Gemini outputs. By default, Gemini only samples or tokenizes one frame each second (of the video playtime). Therefore, we adjusted the video input (original recording 54 fps) to 1, 2, 6, 13, 27, 54 fps which corresponds to a 54, 27, 9, 4.2, 2, and 1 time slowing of frame rates with as shown in Fig. 2C. Using a paired t-test, frames rates of 6 (p <0.01), 13 (p<0.01), 27 (p<0.001), 54 (p<0.01) decreased the accuracy of Gemini classification (Fig. 2C). Although at 2 fps (p = 0.9) did not significantly alter mean accuracy of the model, it increased the variability, i.e. standard deviation, from 0.07 (1fps) to 0.11.

The inter-rater analysis demonstrated that trained (and blinded to outcome) human scorers exhibited similar levels of inter-rater agreement, as measured by Cohen’s kappa, when evaluating mice in the pre-stroke or no-stroke condition (0.693 +/- 0.057), during the early stroke phase (0.831 +/- 0.076), and in the late stroke phase (0.7 +/- 0.047) (Fig. 3A). However, only nine sessions achieved kappa values above 0.80 (substantial agreement), while five sessions fell below the moderate agreement threshold of 0.60. These observations indicate the presence of scorer bias and variability that can influence outcome assessments. When treating Rater 1 (who had the most experience) as the gold standard, Rater 2’s performance remained robust, achieving kappa values greater than 0.80 across all conditions, as shown in Fig. 3A. Given these findings on agreement and accuracy, all the following results are compared to Rater 1 as gold standard. Kappa Values were also calculated for differences between Gemini and human Rater 1. Although a lower average Kappa value of 0.43 +/- 0.24 was found between Gemini and the Rater 1, this Kappa value indicates there is still an overall fair agreement. Notably, 6/16 (pre-stroke or no stroke condition animals) were in strong agreement (Fig S2).

**Figure 3.**
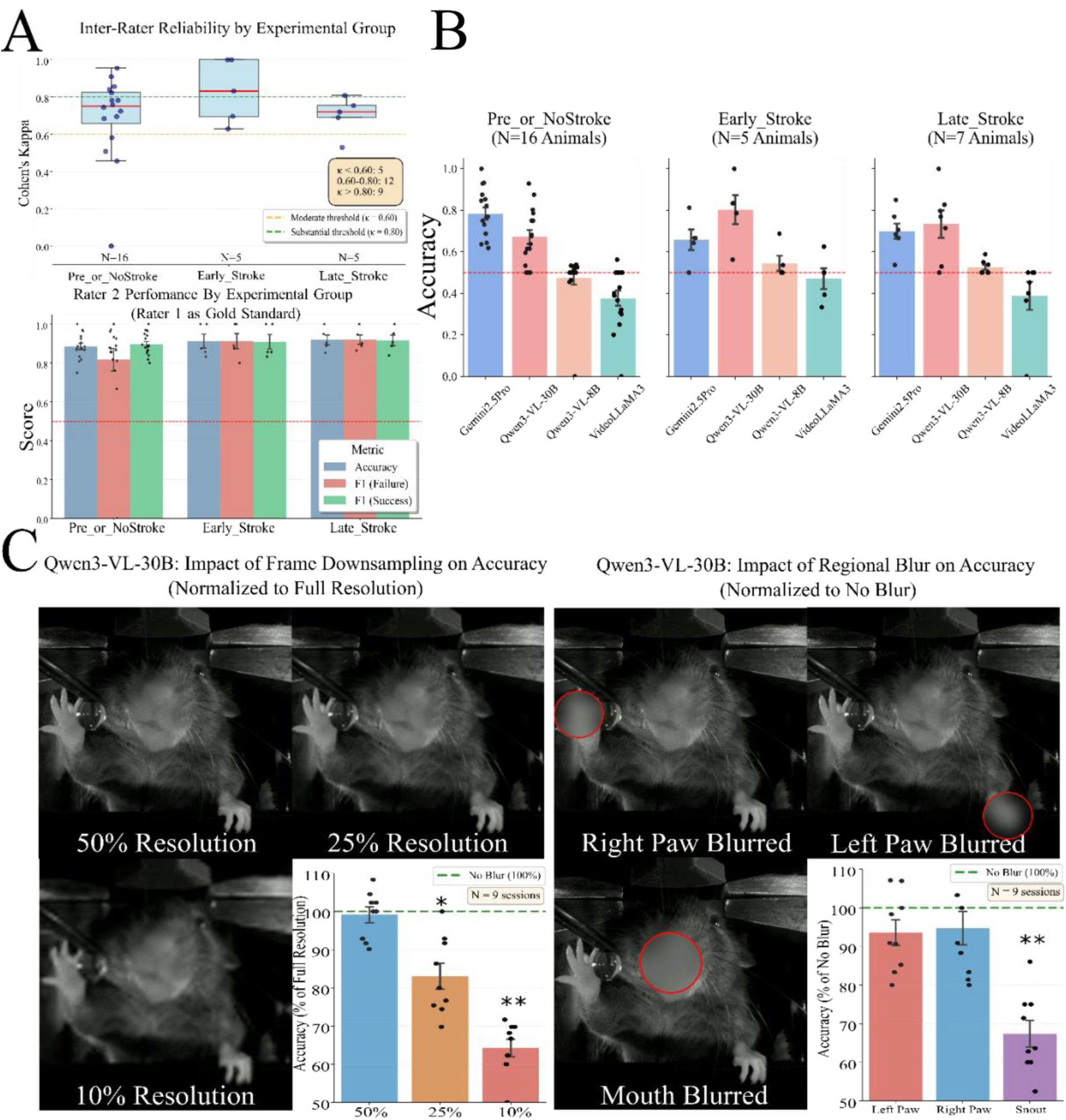
Model Performance on Water Reaching Task. **A.** Inter-rater reliability, as measured by Cohen’s kappa, between two expert human scorers across different experimental groups: Pre-or-No-Stroke, Early Stroke, and Late Stroke. The dashed lines indicate moderate (κ = 0.60) and substantial (κ = 0.80) agreement thresholds and number of sessions that passed the corresponding threshold. **B.** Accuracy of each tested model on classifying water reaching trials across experimental groups. Data is calculated per session (N) which consists of multiple trial videos. **C.** The effect of video resolution and part specific blurring on model performance. The left panel shows example frames at 50%, 25%, and 10% of the original resolution. The bar chart on the right shows the corresponding drop in classification accuracy compared to the original, un-blurred video (green dashed line). Each point represents a session of 6–88 trials; the ground truth is expert hand-scored results. Data are mean ± SEM. B. Comparison of Gemini and human accuracy after corrections. Data are mean ± std. C. The blurred region is enclosed by a red circle. *p < 0.05, **p < 0.01, and ***p < 0.001. Each individual point represents a session composed of 6-88 trials.

Performance of the four video LLM models on the accuracy (versus a Rater 1 as gold-standard) of scoring water-reaching success of mice performing across pre-stroke, early-stroke, and late-stroke animal groups are shown in Fig 3B. A two-way ANOVA (model x group) revealed a highly significant overall difference in classification accuracy among the models (F = 1.795627, p < 0.001). Post-hoc analysis with Tukey’s HSD correction identified: Qwen3-VL-30B (0.71 +/- 0.15) and Gemini 2.5 Pro (0.67 +/- 0.18) demonstrated higher (p<0.001) accuracy compared to Qwen-3-VL-8B (0.50 +/- 0.11) and VideoLLamma3 (0.40 +/- 0.15). Further post-hoc analysis revealed that Qwen-3VL-30B had a higher accuracy compared to Gemini 2.5 Pro in evaluating success and failure in mice during early stroke phases. Although some differences between mean accuracy measures Gemini 2.5 Pro and Qwen3-VL-30B in different treatment groups, no statistical significance was found (model x group interaction effect: p = 0.12). Taken together, these statistical results identify Qwen-3VL-30B and Gemini 2.5 Pro as the only accurate models tested for the automated analysis of this behavioral task. Further looking into the specific classification, F1 scores for failure and success classification seem to be more imbalanced in Qwen3-VL-8B and VideoLLamma3 than Qwen3-VL-30B and Gemini 2.5 Pro (Fig S3) and may explain the differences observed in accuracy measures. Considering cost of analysis (approximately 0.5 USD per video) and given similar performance, further analyses were then conducted with the Qwen-3VL-30B model.

Next, the importance of visual information for trial classification was evaluated in the Qwen-3VL-30B model. Nine sessions with the highest baseline accuracy were selected, and their trial video clips were systematically altered in resolution or feature blurring as described below. To reduce effective resolution while controlling total token input to the model, videos were first down sampled and then resized to their original resolution through bilinear interpolation. Reducing the resolution to 25% (mean accuracy = 83.0%, *p* = 0.011) and 10% (accuracy mean = 64.3%, *p* = 0.004) of the original resulted in a significant, resolution-dependent decrease in model accuracy (Fig. 3C left), as determined by the Wilcoxon signed-rank test. Using these blurring procedures we estimated that pixel sizes greater than 0.28 mm led to decreases in accuracy.

The effect of selectively blurring key body regions was also examined on accuracy. Blurring the right paw (mean = 94.8%, *p* = 0.20) or left paw (mean = 93.6%, *p* = 0.21) did not produce statistically significant changes in accuracy. In contrast, blurring the snout region significantly impaired model accuracy (mean = 67.4%, *p* = 0.039), indicating that orofacial cues are critical for discriminating successful versus failed trials, results are illustrated in Fig 3C right.

Given that the capabilities of video LLMs can extend well beyond simple classification (32), we examined whether they can provide rich, frame-level descriptors of behavior without fine-tuning. Due to accuracy being highest in no stroke or pre-stroke animals for Gemini 2.5 Pro, we further analyzed additional outputs from the prompt for all trial videos (n = 1,058) in these animals. First, Gemini was asked to provide answers for the following descriptors: (1) tongue_contact (whether the tongue touched the water reward), (2) water_drop_stable (whether the water drop remained stable after delivery), (3) water_spilled (whether water spilled from the paw during the task), (4) percentage_consumed (visual estimate of how much of the water drop was consumed), and (5) timeline of events (frame-by-frame event classification and descriptions). These outputs were manually evaluated and curated by an expert (with access to all outputs of the video from Gemini 2.5 Pro) and deemed accurate only when the scorer agreed with the model’s result on the video evaluated. An illustration of these outputs is shown in Figure 4A, B.

**Figure 4.**
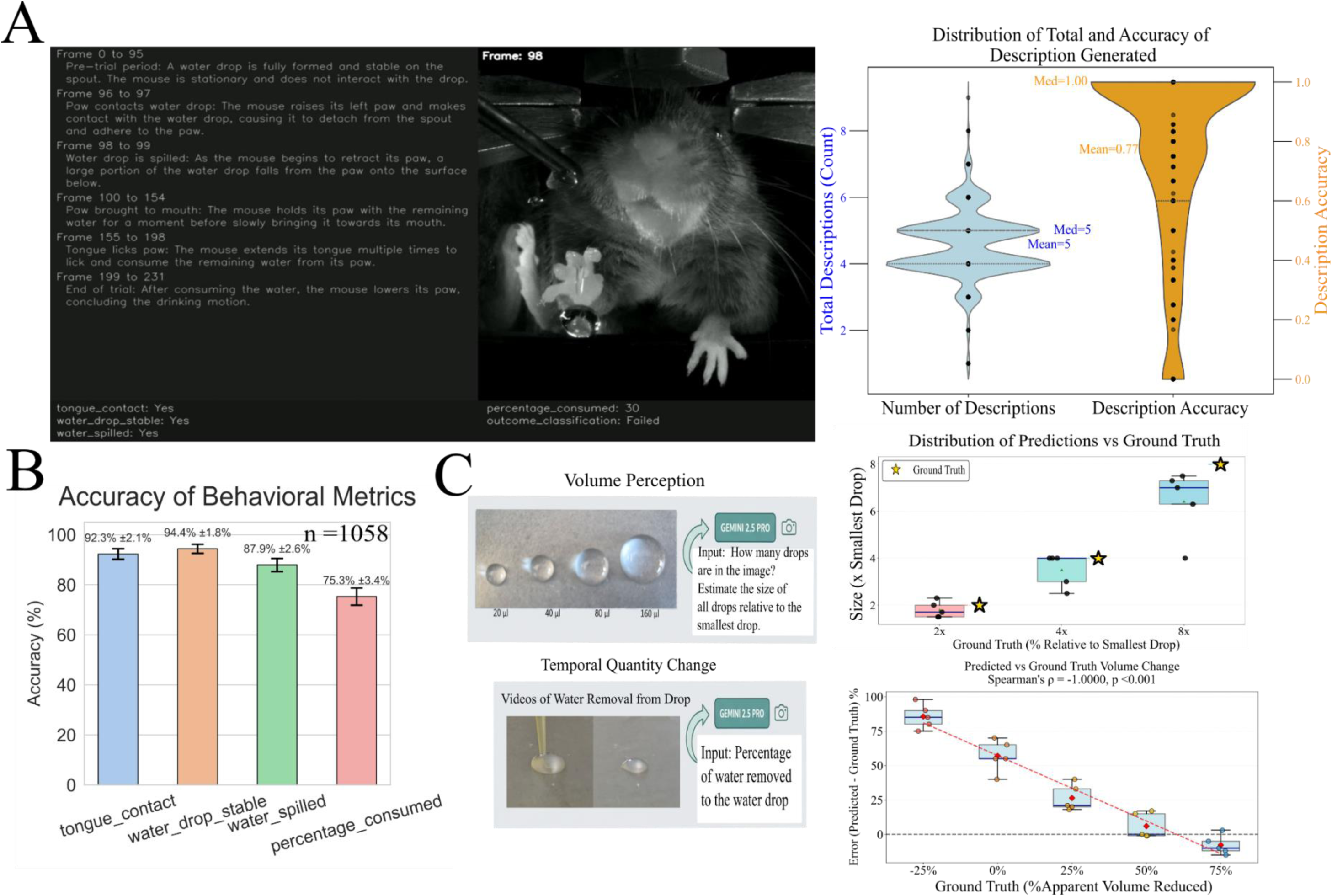
Gemini 2.5 PRO Analysis Performance on Head-Fixed Mouse Water-Reaching Task and Water Drop Quantification. **A.** Example of behavior segmentation by Gemini 2.5 PRO. Comparison of Gemini and human accuracy after corrections. Data are mean ± SEM. **B.** Gemini performance in video segmentation (frame classification by action/content). Data are mean ± 99% CI. The red line shows random (chance-level) accuracy. **C.** Evaluation of Gemini to quantitatively detect water drop sizes in still images and volume changes in videos. Images of water drops were presented and the model estimated size compared to the smallest drop (top graph). Videos showing varying percentages of volume removal from a 80µl water drop were presented to evaluate the model’s ability to track changes in water drop size (bottom graph). In this case large volume reductions simulating consumption of water by the mouse were reported with lower error.

Importantly, Gemini also provides temporal localization of events with per second resolution. As stated, before the temporal resolution of behavior events can be further enhanced by manipulating the sampling rate in frames as the default for Gemini is to sample at 1 Hz by skipping frames. The model can be used in a higher temporal resolution mode by re-sampling the time base: for example, if frames were collected at 54 Hz for 1 s, the time per frame can be reset to create 54 s of 1 Hz data so that all frames will used. As illustrated in Figure 4A, Gemini can detect brief events, such as water spillage lasting only 1–2 frames. Also shown in Figure 4A, Gemini generates on average five descriptions per video (median = 5). A deeper analysis revealed that the average per-video accuracy or in agreement with the evaluator of these descriptions (correct descriptions / total descriptions) was 0.77, with a median of 1.0; notably, 549 of 1,058 videos contained entirely correct descriptions. All 1058 videos are available at the current link (https://osf.io/24euy). For the descriptions tongue_contact, water_drop_stable, and water_spilled, Gemini achieved high accuracies of 92.3 ± 2.1%, 94.4 ± 1.8%, and 87.9 ± 2.6%, respectively, indicating these events were detected reliably and consistently. In contrast, percentage_consumed showed lower agreement with expert scoring, likely because Gemini either may not perceive quantitative amounts or cannot relate visual changes to quantitative measurements. This suggests that Gemini captures key instructed behavioral events robustly but has more difficulty with fine-grained quantitative judgments.

To evaluate the Gemini ‘s quantitative abilities, it was presented with images of four water drops of increasing volumes (20 μl, 40 μl, 80 μl, and 160 μl) and tasked with determining their sizes relative to the smallest drop (Fig. 4C). While the model was able to correctly detect the number of drops in all test cases (N = 5), it was not precise in measuring the exact relative volumes, showing a tendency to underestimate. However, the model demonstrated a clear ability to discern relative sizes, as its predictions showed a significant positive linear correlation with the actual volume (Pearson’s r = 0.57, p=0.03). When presented with videos depicting volume reduction in an 80μl water drop, the model struggled to distinguish small percentage changes. Interestingly, while Gemini struggled to predict small additions or removals (+/- 25% volume), its error rate for removals decreased linearly (Pearson’s r = -0.94, p < 0.01) as the amount of water removed increased indicating lower error near the 70% consumption threshold used to score videos.

## Discussion

Interestingly, trained human scorers can reliably classify trial outcomes without 3D reconstruction or multi-view recordings as we have shown; instead, they rely on a small set of salient visual cues that are often stable across camera setups. This observation suggests that accurate behavior scoring depends less on full kinematic reconstruction and more on extracting task-specific, visually unmistakable events. In this context, the current study provides a comprehensive evaluation of three prominent large language models for automated classification of trial outcomes in a head-fixed mouse water-reaching task. By directly comparing their performance to expert human annotations, we assess whether video-language models can capture the essential visual information needed for determining task success, despite the absence of explicit pose tracking or depth information. Results demonstrated that Qwen-3-VL-30B and Gemini 2.5 Pro were the strongest performers and suggest prompt engineering techniques can influence quality of outputs. Although Gemini 2.5 Pro achieved the highest overall accuracy, its performance was not statistically superior to Qwen3-VL-30B when all experimental groups were combined, although see discussion of stroke performance below. Notably within the Gemini 2.5 Pro results, accuracy in the no/pre-stroke condition was trended higher than in the early-stroke and later-stroke condition. Classification performance—F1-score—for the success category remained superior to that for failure across all sessions and conditions for the Gemini 2.5 Pro model. This likely reflects the fact that successful water consumption presents clear, consistent, and robust visual cues (e.g., paw movement, tongue contact, and water uptake), whereas failed attempts are more heterogeneous and less visually well-defined. Interestingly, Qwen-3-VL-30B exhibited relatively better performance in disease-relevant conditions (early stroke: 0.8+/-0.16; later stroke: 0.73 +/- 0.17; not statistically significant), suggesting enhanced sensitivity to altered motor behaviors and increased utility in studies of neurological impairment.

In contrast, the smaller Qwen3-VL-8B and the VideoLLaMA3 (a 7B model) models constituted a significantly lower performance tier. Their failure was not merely a matter of lower accuracy but a profound classification bias, evident from their near-zero F1 scores for “Success” trials (Fig S3). These models effectively learned to default to a “Failure” classification, rendering them practically useless for an analysis that requires the correct identification of both outcomes even after. To avoid bias due to class imbalance, we subsampled the dataset to achieve equal representation of successful and failed trials during evaluation. The performance gap between the top and bottom models was not only statistically significant (p < 0.001). This suggests that a certain level of model complexity and parameter size is necessary to capture and describe the visual details of complex motor behaviors.

To investigate the effects of quality of input, we tested the reliance of Qwen3-VL-30B on specific visual information. Our findings reveal that the model’s accuracy is highly dependent on image fidelity, with performance significantly degrading as effective video resolution was reduced to 25% and 10% of the original through bilinear interpolation. This indicates the importance of fine-grained visual details rather than just gross movements for video LLMs. At the original resolution, the imaging scale was 0.14 mm per pixel. This suggests that the minimal features required for accurate classification become degraded once the effective resolution drops to between 0.28 and 0.57 mm per pixel corresponding to 50% and 25% effective resolution, respectively. Moreover, selective blurring experiments revealed that orofacial cues are the most critical feature for classification. While obscuring either paw (so it became an un-recognizable blur) did not significantly impact performance, blurring the facial region led to a significant drop in accuracy. This strongly suggests that the model has learned to identify the most biologically relevant endpoint of the task—the successful delivery of the water droplet to the mouth—mimicking the analytical process of a human expert.

Most impressively, the ability of Gemini 2.5 Pro to generate descriptive text that segments behaviors temporally offer a transformative opportunity for accelerating research. Even though not all descriptions generated are correct, a substantial fraction are and can be curated or confirmed to rapidly generate large amounts of high-quality training data. This process addresses a key challenge in developing video understanding systems: the creation of extensive, accurately labelled datasets for fine-tuning large language models (LLMs). The outputs could also support the training or refinement of other neural network–based approaches, such as A-SOiD (14), serving as a powerful first pass or complementary tool to conventional labelling platforms like BORIS (33). The ability of Gemini to perform complex segmentation tasks so effectively, where other tested models failed, is suspected to be due to its greater number of model parameters and assumed complexity. While the model remains proprietary without a supporting publication on the architecture, the observed capabilities suggest an exceptionally large parameter count, likely exceeding 200 billion, which may facilitate advanced video understanding abilities.

Given the rising spatial-temporal understanding capabilities of recent video LLMs (34,35), we further tested specifically the ability of Gemini on estimating the volume of a water drop, an object of interest in our task. Judging the relative size of an object within a single static image was effective, as evidenced by the model’s robust detection of drop numbers and a significant positive linear correlation of predicted size with actual volumes in the water drop. Unfortunately, the model’s ability to precisely quantify temporal volumetric changes (such as water being sucked from a drop over a period of time) was more limited. Accurate detection of dynamic alterations across time appears to require robust visual alterations in the drop meaning that gradual changes may go un-noticed by a video LLM. This is supported by the observed linear decrease in error for predicted percentage removed as the true amount of water removed increased. Therefore, for applications requiring the tracking of evolving amounts, enhanced visual fidelity or more pronounced physical transformations are critical for the model to perform reliably. Perhaps, through adjustment of prompts the model could be encouraged to spot volume change over slower timescales.

The presented approach can substantially lower the technical barrier for behavioral neuroscience by enabling automated trial outcome classification without the need for specialized imaging hardware or sensor systems. This framework scales readily and can facilitate standardized, reproducible analyses across tasks and laboratories, opening the door to high-throughput behavioral pipelines while reducing labor-intensive manual annotation and minimizing human bias. As large video-capable language models continue to advance rapidly, they hold the potential to transform behavioral neuroscience in a manner analogous to the impact of convolutional neural networks on image analysis, accelerating discovery and allowing researchers to focus on interpretation rather than scoring.

Most high-performing models require significant computational resources and are not broadly accessible in typical biological laboratories. Furthermore, Full training is impractical, demanding millions of video–text pairs and data-centre GPUs with over 300 GB of VRAM (36). Fortunately, open-source models such as Qwen enable domain adaptation through LoRA, which updates small low-rank adapter layers while keeping most parameters frozen significantly reduce GPU memory requirements. However, thousands of high-quality video–text pairs are still required and is a highly manual process (2,37,38). In disease-relevant contexts such as stroke or neurodegenerative models, fine-tuning model parameters is likely necessary to capture altered kinematics and behavioral variability. As previously mentioned, Gemini’s accessibility further supports this process by facilitating the generation of high-quality image and video–text pairs. Interestingly, the strong performance of general-purpose video LLMs suggests that training on large-scale human-action datasets may allow cross-domain transfer to rodent behaviors, potentially complemented by incidental exposure to animal footage during pretraining (25,39); however, systematic evaluation is needed to confirm this possibility. Future work leveraging additional open-source models and diverse behavioral assays will be critical for assessing generalizability and determining how broadly capabilities of video LLMs extend beyond the current task.

## Disclosures

The authors declare that there are no financial interests, commercial affiliations, or other potential conflicts of interest that could have influenced the objectivity of this research or the writing of this paper.

## Acknowledgements

Pumin Wang and Cindy Jiang for surgical assistance. Jeffrey M LeDue and Federico Bolanos for technical assistance. This work was supported by a Canadian Institutes of Health Research (CIHR) project grant PJT-180631 to T.H.M. THM was also supported by the Brain Canada Neurophotonics Platform, a Natural Science and Engineering Council of Canada (NSERC; GPIN-2022-03723). This work was supported by resources made available through the Dynamic Brain Circuits cluster and the NeuroImaging and NeuroComputation Centre at the UBC Djavad Mowafaghian Centre for Brain Health (RRID SCR_019086) and made use of the DataBinge forum.

## Code and Data Availability

All data and relevant scripts are available at the following open science framework and GitHub repositories https://osf.io/24euy and https://github.com/tf4ong/videollm, respectively.

## Notes

https://github.com/tf4ong/videollm

https://osf.io/24euy

### Competing Interest Statement

The authors have declared no competing interest.

