## Supplemental Figures for "Towards Video-LLM Driven Workflow for Behavioral Segmentation and Scoring in Mice Performing a Skilled Water Reaching Task: An Evaluation of Recent LLM Models"

### Supplemental Information

#### • System Instruction

- You are a research assistant tasked with objectively scoring a mouse's behavior in a water-reaching trial.
- Your analysis must be based only on clear, indisputable visual evidence. Pay extremely close attention to the following elements:
- 1. The Water Drop: Note its formation, its position on the spout, and any change to its shape or location.
- 2. The Mouse's Tongue (very obvious and similar to that of a dog's): Look for the moment the tongue is extended or if the tongue physically touches the water.
- 3. The Mouse's Paws: Observe if the paws lift off the surface and move towards the spout. Mainly the mouse's right paw (the mouse's perspective).
- Most importantly, the mouse is head fixed meaning it cannot move its head and its tongue cannot lick the spout directly (the mouse's perspective and the one closest to the metal spout when the paws are at rest).
- The water drop can only be consumed by its right paw reaching the water drop on the spout and bringing it close to its mouth for it to lick.
- If the mouse's right paw did not come near to the water drop or move at all, it couldn't have drank any of the water drop.
- The water drop is always delivered within the first 30–40 seconds of the video. In some cases, the water drop might not stick to the spout for the mouse to touch.
- Do not infer success or failure; only report the physical interactions you see. In each video, only one drop is delivered.
- If no change or movement (right paw) is detected for a specific feature (e.g., paw, tongue), you must state: "No change was observed."
- Do not repeat the question in your responses.
- Do not describe an event unless there is undeniable visual evidence.

Fig S1. Full system instructions. System instructions were inputted as parameters to Gemini. In other models, system instructions were inputted as past chat history prior to video analysis.

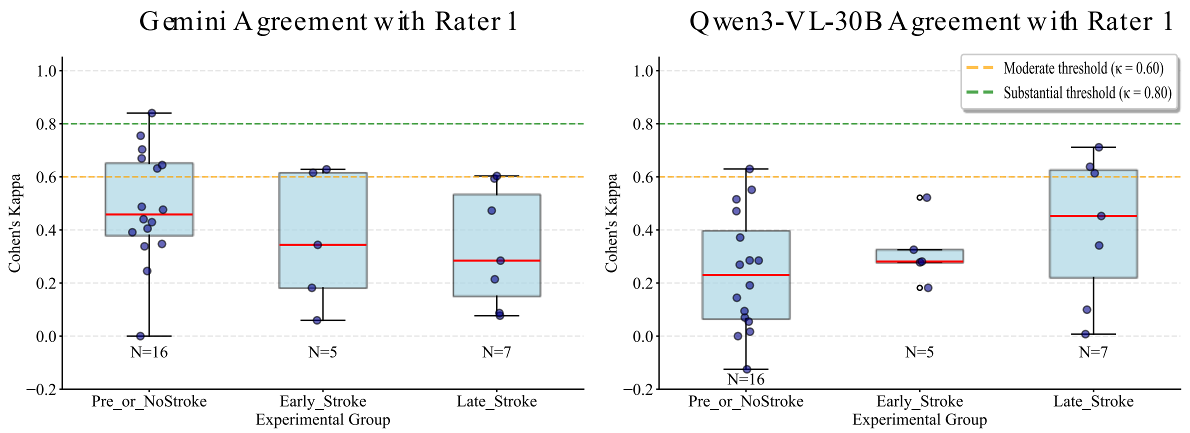


Fig S2. Inter-rater agreement between Gemini and Qwen3-VL-30B with Rater 1. Inter-rater reliability is measured Cohen's kappa across different experimental groups: Pre-or-No-Stroke, Early Stroke, and Late Stroke. The dashed lines indicate moderate (κ = 0.60) and substantial (κ = 0.80) agreement thresholds and number of sessions that passed the corresponding threshold.

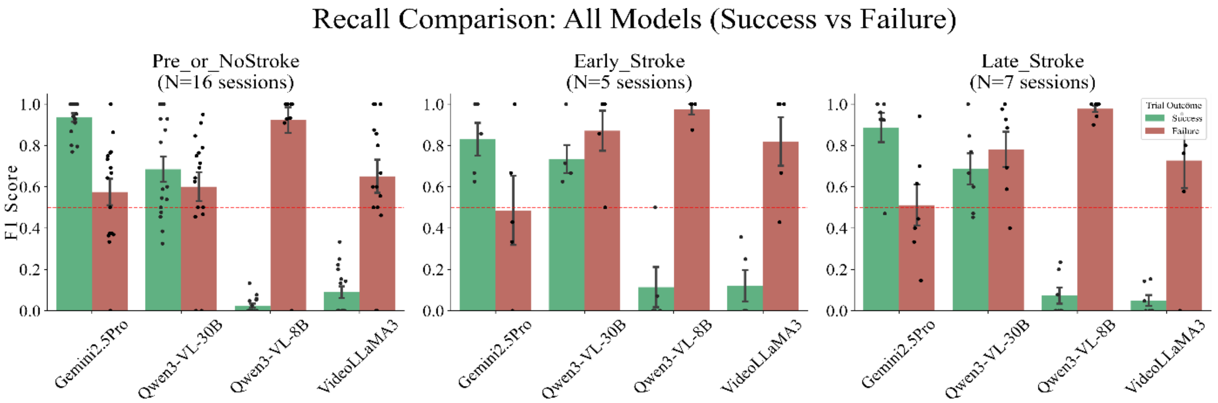


Fig S3. F1 score metrics for success and failure across video LLM models. F1 score for four different models (Gemini 2.5 Pro, Qwen3-VL-30B, Qwen3VL-8B, and VideoLLaMA 3) in classifying trial outcomes as either success and failure. The performance is evaluated across three distinct experimental groups: Pre-or-No-Stroke (N=16 sessions), Early Stroke (N=5 sessions), and Late Stroke (N=7 sessions). Data expressed as mean +/- std.
